# bMAE: Masked Autoencoder Latent Representations for Bulk RNA-seq Tissues

**DOI:** 10.64898/2026.03.03.709470

**Authors:** Zikang Wan, Marko Zolo Gozano Untalan, Danilo Vasconcellos Vargas

## Abstract

Bulk tissue RNA-sequencing data from large-scale consortia such as GTEx provide comprehensive gene expression profiles across diverse human tissues. However, the high-dimensional nature of bulk RNA-seq data, combined with technical noise and batch effects, poses challenges for downstream analyses. While dimensionality reduction methods are routinely applied, standard approaches such as PCA often fail to optimally preserve tissue-discriminative information and exhibit poor generalization to unseen tissue types. We developed a masked autoencoder for bulk tissue RNA-seq that learns compressed latent representations through self-supervised learning with variable masking schedules. Evaluated on GTEx data (31 tissue types, 19,788 samples, 19,308 genes), our method substantially outperformed all baselines in leave-one-tissue-out (LOTO) cross-validation. We achieved mean silhouette 0.20, ARI 0.58, and NMI 0.84 versus best baseline UMAP (0.007, 0.25, 0.47), representing 28.6-fold, 2.3-fold, and 1.8-fold improvements. Remarkably, within held-out tissue categories containing subtissues, our latent space revealed hierarchical structure with enhanced subtissue separation (silhouette 0.16, ARI 0.35, NMI 0.36) versus baselines (silhouette 0.15, ARI 0.20, NMI 0.20), despite never observing these distinctions during training. The method compressed 19,308 genes to 128 dimensions while preserving multi-scale structure.

## Introduction

### Bulk Tissue RNA-seq Gene Expression

Bulk tissue RNA-sequencing has emerged as a foundational technology for characterizing gene expression across diverse biological contexts. Large-scale consortia such as the Genotype-Tissue Expression (GTEx) project have generated comprehensive transcriptomic profiles, with the Adult GTEx release containing approximately 19,788 samples from 54 tissue sites across 946 donors[3]. These data provide unprecedented opportunities to understand tissue-specific gene regulation[3, 1], identify disease biomarkers[7, 16], and help understand fundamental biological processes[13, 12]. However, the high-dimensional nature of bulk RNA-seq data, with expression measurements for tens of thousands of genes per sample, poses significant computational and analytical challenges for downstream analyses including tissue classification, differential expression analysis, and data integration[8, 4, 21].

### Dimension Reduction

Traditional dimensionality reduction approaches, particularly principal component analysis (PCA), are routinely applied to mitigate the curse of dimensionality in RNA-seq data[19, 20]. While PCA effectively reduces data complexity through linear transformations, it often fails to optimally preserve nonlinear biological relationships and tissue-discriminative information in reduced representations. Moreover, standard dimensionality reduction methods exhibit poor generalization when applied to unseen tissue types, limiting their utility in transfer learning scenarios. Despite these limitations, bulk RNA-seq maintains irreplaceable advantages over single-cell technologies, including substantially lower per-sample costs, higher gene coverage with minimal sparsity, greater accessibility to fixed specimens from clinical archives, and the ability to capture population-level tissue signals that reflect true biological composition rather than sampling artifacts.

### Masked Autoencoders

Recent advances in self-supervised learning have demonstrated remarkable success in learning robust representations from high-dimensional biological data. Inspired by masked language modeling in natural language processing (BERT)[9] and masked autoencoders in computer vision (MAE)[14], several methods have applied masking strategies to single-cell RNA-seq data. The scMAE framework introduces partial corruption to gene expression profiles and leverages masking prediction to capture gene-gene correlations[10], while scDECL employs contrastive learning with masking as a pretext task for clustering enhancement[11]. These approaches demonstrate that forcing models to reconstruct randomly masked values encourages learning of biologically meaningful patterns rather than memorizing noise or technical artifacts. The success of masked autoencoders in single-cell contexts suggests potential for similar gains in bulk tissue analysis.

Despite extensive bulk RNA-seq resources accumulated over two decades covering virtually all human tissue and organ types, masked autoencoding methods remain underexplored for bulk tissue transcriptomics. The distinct characteristics of bulk data, including higher signal-to-noise ratios, reduced sparsity, and population-level averaging effects, suggest that techniques optimized for single-cell data may not directly transfer.

In this work, we demonstrate that masked autoencoders with variable masking rate schedules can learn generalizable tissue representations from bulk RNA-seq data that significantly outperform raw expression profiles, PCA-reduced representations and UMAP-reduced representations. Through rigorous leave-one-tissue-out cross-validation on the GTEx consortium dataset, we show that our approach captures transferable biological principles enabling accurate classification of entirely unseen tissue types, addressing a critical gap in representation learning for bulk transcriptomics.

## Dataset and Problem Formulation

### Adult GTEx

We utilized bulk tissue RNA-seq data from the Genotype-Tissue Expression (GTEx) project, which represents a comprehensive publicly available resource for studying gene expression across diverse human tissues. The data used for this research were obtained the GTEx Portal on December 1, 2025. The Adult GTEx dataset comprises 946 postmortem donors and 19,788 samples spanning 31 major tissue types. Data were collected using standardized protocols to minimize technical variation, including consistent postmortem intervals, tissue procurement procedures, and RNA extraction methods. All samples were sequenced using Illumina platforms with comparable sequencing depths to ensure data quality and comparability across tissues.

Gene expression values were expressed as Transcripts Per Million (TPM), which normalizes for both gene length and total sequencing depth, enabling fair comparisons across genes and samples. We restricted our analysis to 19,308 protein-coding genes. Each sample is represented as a TPM vector **X** ∈ ℝ^*G*^, where *G* denotes the number of genes. The complete dataset forms a gene expression matrix **X** ∈ ℝ^*N*×*G*^, where *N* is the total number of samples. Tissue labels are available for all samples, enabling supervised evaluation of learned representations; however, these labels were not used during model training, maintaining the self-supervised nature of our approach. The model receives and outputs values transformed via *r*(tpm) = log(tpm + 1)/ log(1 × 10^6^ + 1), but all values in metrics are calculated in the original TPM space via the inverse *r*^−1^.

### Masking and Recovery

The core of our self-supervised learning approach is a masking and recovery framework that forces the model to learn robust gene expression patterns by reconstructing randomly corrupted inputs. Given a gene expression vector **X** ∈ ℝ^*G*^ representing a single bulk tissue sample, we define the masking operation as follows. We first sample a binary masking indicator vector ***m*** ∈ {0, 1}^*G*^ where each element *m*_*j*_ is drawn independently from a Bernoulli distribution with masking probability *p*_mask_: *m*_*j*_ ~ Bernoulli(*p*_mask_). The masked input is then constructed as **X**_masked_ = **X** ⊙ (**1** − ***m***), where ⊙ denotes element-wise multiplication and **1** is a vector of ones. This operation sets the expression values of masked genes to zero: *x*_masked,*j*_ = 0 if *m*_*j*_ = 1, and *x*_masked,*j*_ = *x*_*j*_ otherwise.

The recovery objective requires the model to predict the original expression values from the corrupted input. Formally, we seek to learn a function *f*_𝜃_ : ℝ^*G*^ → ℝ^*G*^ parameterized by neural network weights 𝜃 such that *f*_𝜃_(**X**_masked_) ≈ **X**. This reconstruction task differs fundamentally from standard denoising objectives: rather than adding Gaussian noise, masking creates discrete information gaps that require the model to leverage learned gene-gene correlations and biological constraints to infer missing values. The stochastic nature of masking ensures that the model cannot simply learn an identity mapping and must instead capture robust statistical patterns in gene expression that generalize across different masking configurations. Importantly, only the masked positions contribute to the training loss (detailed in Section 3.3), preventing the model from trivially copying unmasked values and ensuring that learning signal derives exclusively from reconstruction of corrupted information.

To quantitatively assess reconstruction quality, we employ two complementary metrics. First, we compute Pearson correlation coefficient 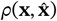 between true and reconstructed gene expression vectors and the Spearman correlation coefficient 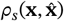, which measures preservation of global expression patterns independent of scale.

### Leave One Tissue Out

To rigorously evaluate the generalization capability of learned representations, we employed a leave-one-tissue-out (LOTO) cross-validation strategy that tests the model’s ability to classify samples from entirely unseen tissue types. The GTEx dataset contains numerous sub-tissues within broader tissue categories. For example, the Adipose tissue includes Adipose - Visceral (Omentum) and Adipose - Subcutaneous. We obtained the major tissue categorization simply by designating the name to left of the dash(-) to be the major tissue classification and to the right to be the subtissue classification. This grouping ensures that our evaluation tests generalization at a biologically meaningful level of tissue identity rather than subtle regional variations. This resulted to 10 major tissues.

The LOTO procedure operates as follows. For each major tissue type *t* ∈ {1, …, *T*}, we partition the dataset into a training set containing all samples from tissues {1, …, *T*}\ {*t*}and a test set comprising all samples from the held-out tissue *t*. We train a masked autoencoder from scratch on the training set using the self-supervised masking objective described in Section 2.2. After training for a set number of epochs (detailed in Section 3.4, we extract latent representations for all test samples from tissue *t* and perform clustering analysis in this learned latent space. We evaluate clustering quality using Adjusted Rand Index (ARI) and Normalized Mutual Information (NMI) by comparing predicted clusters to ground-truth tissue labels. This process is repeated for each of the *T* tissue types, yielding *T* independent measurements of generalization performance. Final metrics are reported as mean ± standard deviation across all LOTO folds.

The LOTO evaluation differs critically from standard *k*-fold cross-validation, where samples are randomly split without regard to tissue identity. Random splitting allows the model to observe examples from all tissue types during training, testing only whether it can generalize to new samples from known tissues which is a substantially easier task. In contrast, LOTO presents a domain generalization challenge: the model must learn tissue-discriminative features from a subset of tissue types and apply them to classify samples from tissue types never observed during training. This strict evaluation protocol tests whether learned representations capture fundamental, transferable principles of tissue identity or merely memorize tissue-specific patterns in the training data. Success in LOTO validation indicates that the latent space encodes biologically general axes of variation: such as epithelial versus mesenchymal characteristics, metabolic activity profiles, or developmental lineage markers that enable zero-shot recognition of novel tissue contexts.

The biological significance of LOTO generalization extends beyond methodological rigor to practical applicability. In real-world scenarios, researchers frequently encounter rare tissue types, novel disease contexts, or specialized anatomical regions with limited available samples. A representation learning method that successfully generalizes in LOTO validation demonstrates potential for few-shot learning applications, where accurate classification can be achieved with minimal labeled examples from new tissue types. Furthermore, strong LOTO performance suggests that the model has learned interpretable biological features rather than exploiting dataset-specific artifacts or batch effects that would not transfer to held-out tissues. This generalization capability is essential for enabling transfer learning across diverse bulk RNA-seq datasets from different laboratories, sequencing platforms, and biological conditions.

## Modeling

### Architecture

Our masked autoencoder employs a symmetric multilayer perceptron (MLP) architecture with separate encoder and decoder pathways. The encoder progressively compresses gene expression vectors through three hidden layers with dimensions *G* → 2048 → 1024 → 512 → 128 → *d*, where *G* = 19, 308 is the number of input genes and *d* = 128 is the latent bottleneck dimension. Each hidden layer is followed by LayerNorm [5], GELU [15](Gaussian Error Linear Unit) activation, and dropout (rate = 0.1). The decoder mirrors this architecture in reverse: *d* → 128 → 512 → 1024 → 2048 → *G*, with identical normalization, activation, and dropout[18] after each layer. The final output layer applies a sigmoid activation to constrain reconstructed values to [0,1], matching the normalized input range.

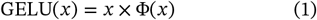

where Φ(*x*) is the cumulative distribution function for Gaussian distribution.

We selected GELU activation over ReLU[2] for its smooth, non-monotonic properties that have shown superior performance in representation learning tasks. LayerNorm stabilizes training by normalizing activations within each layer, while dropout provides regularization to prevent overfitting. The symmetric encoder-decoder design with progressively expanding/contracting dimensions creates an information bottleneck at the latent layer, forcing the model to learn compressed representations that capture essential gene expression patterns.

### Mask Scheduling

A key innovation in our approach is the use of a variable masking rate schedule during training, in contrast to the fixed masking rates employed by most prior work. Standard masked autoencoder methods maintain a constant masking probability *p*_mask_ throughout training. For example, consistently masking 30% or 50% of genes in ev-ery training iteration. While this approach is straight-forward, we hypothesized that dynamically adjusting the masking rate could improve learned representation quality by implementing a form of curriculum learning.

Our variable masking schedule modulates *p*_mask_ as a function of training progress marked by the epoch *e* via a cosine schedule with a period of 𝜏.

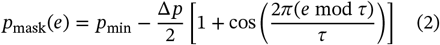

where ∆*p* = *p*_min_ − *p*_max_, and *p*_min_ and *p*_max_ are implementation details in Sec 3.4.

The intuition behind variable masking is that different masking rates encourage learning of different types of features. As illustrated in Figure 2, early in training with high masking rates (e.g., sup *p*_mask_ = 0.9), the model faces a more challenging reconstruction task with limited input information. This difficulty forces the network to identify and leverage robust, high-level patterns in gene expression,such as major tissue-specific expression signatures, co-regulated gene modules, and fundamental biological constraints that are essential for meaningful reconstruction. As training progresses and the masking rate decreases (e.g., inf *p*_mask_ = 0.1), the model can refine these coarse representations by incorporating finer-grained details and gene-specific patterns. This progression mirrors curriculum learning principles, where models first master fundamental concepts before tackling more nuanced distinctions.

**Figure 1:**
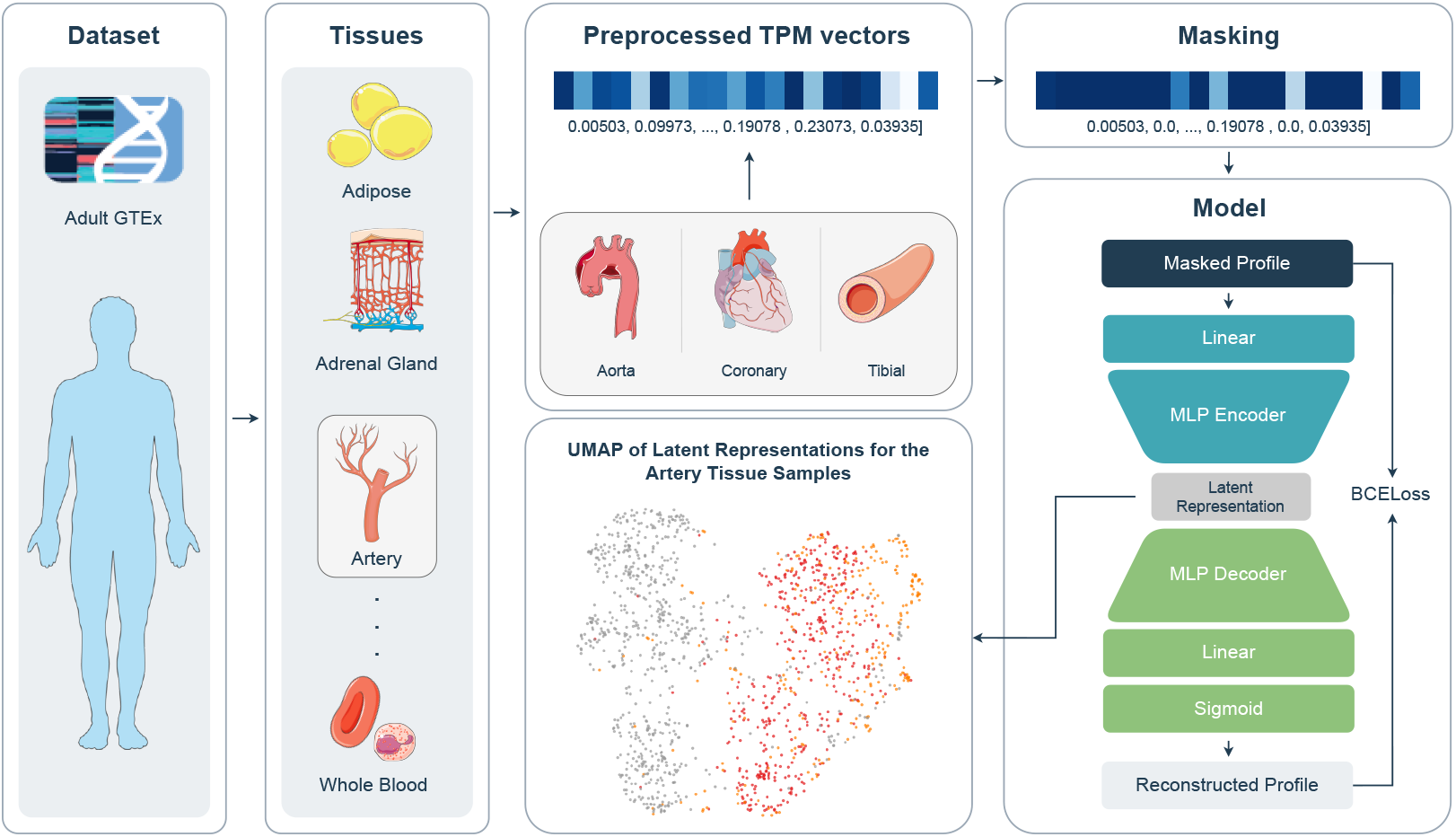
Framework: (1) Dataset is partitioned into major tissues; (2) Subtissues are identified; (3) Profiles are randomly masked according to a schedule; (4) Autoencoder; (5) Latent Space Visualization [Tissues images are adapted from Servier Medical Art (https://smart.servier.com/), licensed under CC BY 4.0 (https://creativecommons.org/licenses/by/4.0/).]

**Figure 2:**
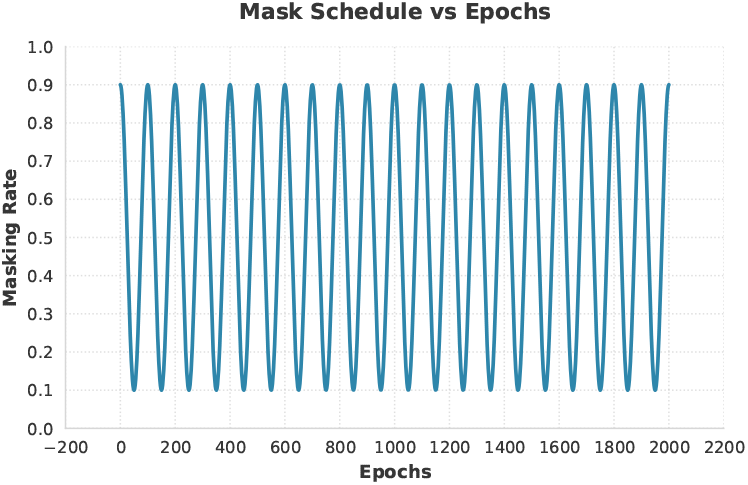
Mask Schedule: Starts on a high masking rate and ends on a low masking rate

From a biological perspective, the variable masking schedule aligns with the hierarchical organization of gene expression in that genes affect each other in cascades which the model would have to use to recover the missing ones [10, 14]. At the highest level, tissues are distinguished by expression of canonical marker genes and activation of tissue-specific transcriptional programs. At finer resolutions, subtissues and cell-type compositions create more subtle expression variations. By starting with aggressive masking that emphasizes learning major axes of variation and gradually reducing masking to capture detailed structure, our schedule encourages the model to build representations that naturally reflect this biological hierarchy.

### Loss Function

We compute binary cross-entropy loss exclusively over masked gene positions. Let ℳ denote the set of masked gene indices. Given normalized expression **X** and reconstruction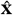:

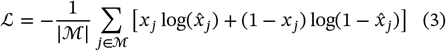

Only masked positions (not unmasked genes) contribute to gradients, forcing the model to learn gene-gene relationships rather than identity mappings.

### Training and Parameters

#### Overview

We implemented the masked autoencoder framework in PyTorch and trained all models on a single NVIDIA GeForce RTX 4090 (24GB VRAM). The model parameters were initialized using the default He initialization. We employed the Adam optimizer[17] with an initial learning rate of 𝛼 = 1 × 10^−4^. Training proceeded for 2000 epochs with a mini-batch size of 128. We employ a *p*_max_ = 0.90 and *p*_min_ = 0.10 for the masking schedule. We use a 𝜏 = 100

#### Shared Configuration across Folds

For leave-one-tissue-out experiments, we trained a separate model from scratch for each fold using only the training tissues using the same hyperparameters on all folds. After training, we extracted latent representations by passing samples through the trained encoder and used these *d*-dimensional embeddings for all downstream clustering analyzes.

## Results and Discussions

### Latent Space

To evaluate the quality of learned representations, we treated each model’s encoder as a dimensionality reduction function *f*: ℝ^19,308^ → ℝ^128^ and systematically compared the resulting latent spaces. We benchmarked bMAE and its ablation variants against established dimensionality reduction methods and raw expression data across multiple clustering quality metrics.

#### Comparison of Methods

We compared 5 approaches for representing gene expression data:

##### Neural network-based method (Ours)

- **bMAE (128 latent dimension)**: Our masked autoencoder with variable masking schedule

##### Classical dimensionality reduction

- **PCA (128 dimensions):** Principal Component Analysis is the most widely adopted linear dimensionality reduction technique in transcriptomics, valued for its computational efficiency, interpretability, and ability to capture the dominant axes of variance in gene expression data. PCA has been extensively used for batch effect correction, quality control, and visualization in large-scale RNA-seq studies including GTEx and TCGA.
- **UMAP (128 dimensions):** Uniform Manifold Approximation and Projection leverages manifold learning and topological data analysis to preserve both local and global structure. Unlike PCA’s linear projections, UMAP constructs a fuzzy topological representation of the high-dimensional data and optimizes a low-dimensional embedding to maintain these topological relationships, making it particularly effective for revealing non-linear patterns and clusters in complex biological datasets[6].

##### Baseline

- **Raw:** Original 19,308-dimensional gene expression profiles without dimensionality reduction

Since bMAE reduces the profiles to 128 dimensions, while PCA and UMAP were configured to match this dimensionality for fair comparison.

#### Clustering Quality Metrics

We assessed latent space quality using three complementary metrics that evaluate different aspects of cluster structure:

##### Silhouette Score

measures how well-separated clusters are by computing the ratio of between-cluster to within-cluster distances for each sample. For sample *i* with within-cluster mean distance *a*(*i*) and nearest-cluster mean distance *b*(*i*):

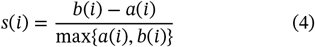

The silhouette score ranges from −1 to 1, where values near 1 indicate samples are well-matched to their own cluster and poorly-matched to neighboring clusters. Negative values suggest potential misclassification. This metric directly quantifies the compactness and separation of tissue clusters in latent space, with higher values indicating clearer tissue boundaries.

##### Adjusted Rand Index (ARI)

quantifies agreement between predicted clusters and true tissue labels by counting sample pairs that are consistently grouped or separated:

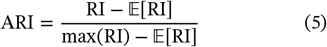

where RI is the Rand Index. ARI is adjusted for chance, yielding values from −1 to 1, where 1 indicates perfect label agreement, 0 represents random clustering, and negative values indicate worse-than-random performance. Unlike silhouette score, ARI explicitly evaluates how well the latent space preserves known tissue identities when subjected to clustering algorithms (we used k-means with *k* equal to the number of tissue types).

##### Normalized Mutual Information (NMI)

measures the mutual dependence between cluster assignments and true labels through information theory:

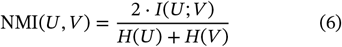

where *I*(*U*; *V*) is mutual information between cluster assignments *U* and true labels *V*, and *H*(⋅) denotes entropy. NMI ranges from 0 (no mutual information) to 1 (perfect correlation), quantifying how much information about tissue identity is retained in cluster assignments. NMI is particularly robust to cluster number imbalances and provides a probabilistic perspective on clustering quality. Similarly, we also used *k*-means to find the clusters with *k* equal to the number of tissue types.

Together, these metrics provide a comprehensive evaluation: silhouette score assesses geometric separation, ARI measures label recovery, and NMI quantifies information preservation.

#### Performance on Complete Tissue Categories

We evaluated all methods using leave-one-tissue-out (LOTO) cross-validation across all 31 tissue types. For each fold, one tissue was held out as the test set while the remaining 30 tissues formed the training set. Neural network models were trained only on the 30 training tissues, while PCA and UMAP were fitted on all of the data, allowing these methods to see the full range of tissues which simulates the usual use of these methods.

Table 1 presents the mean clustering metrics averaged across all 31 LOTO folds. bMAE substantially outperformed all baselines across all metrics, achieving mean silhouette score of 0.1996, ARI of 0.5805, and NMI of 0.8438. Compared to the best classical method (UMAP: silhouette 0.0074, ARI 0.2454, NMI 0.4675), our approach achieved 28.6-fold improvement in silhouette score, 2.3-fold improvement in ARI, and 1.8-fold improvement in NMI.

**Table 1:** Latent space quality metrics averaged across 31 LOTO splits. All methods project to 128 dimensions except Raw. Best results in **bold**, second-best underlined. Values reported as mean ± standard deviation.

| Method | Silhouette ↑ | ARI ↑ | NMI ↑ |
| --- | --- | --- | --- |
| *Neural network-based* |  |  |  |
| bMAE | **0.1996 ± 0.019** | **0.5805 ± 0.036** | **0.8438 ± 0.014** |
| *Classical dimensionality reduction* |  |  |  |
| UMAP | <u>0.0074</u> | <u>0.2454</u> | <u>0.4675</u> |
| PCA | -0.0093 | 0.1913 | 0.3756 |
| *Baseline* |  |  |  |
| Raw | -0.0092 | 0.1814 | 0.3703 |

Among classical methods, UMAP consistently outperformed PCA across all metrics, validating the importance of preserving non-linear structure in gene expression data. However, both classical approaches were substantially inferior to bMAE, suggesting that self-supervised representation learning with reconstruction objectives captures tissue-discriminative features more effectively than unsupervised projections.

Figure 3 shows metric distributions across individ-ual LOTO folds, demonstrating that bMAE’s superiority holds consistently across diverse tissue types. The model achieved the highest scores all 31 folds for all measurements, Silhouette, ARI, and NMI.

**Figure 3:**
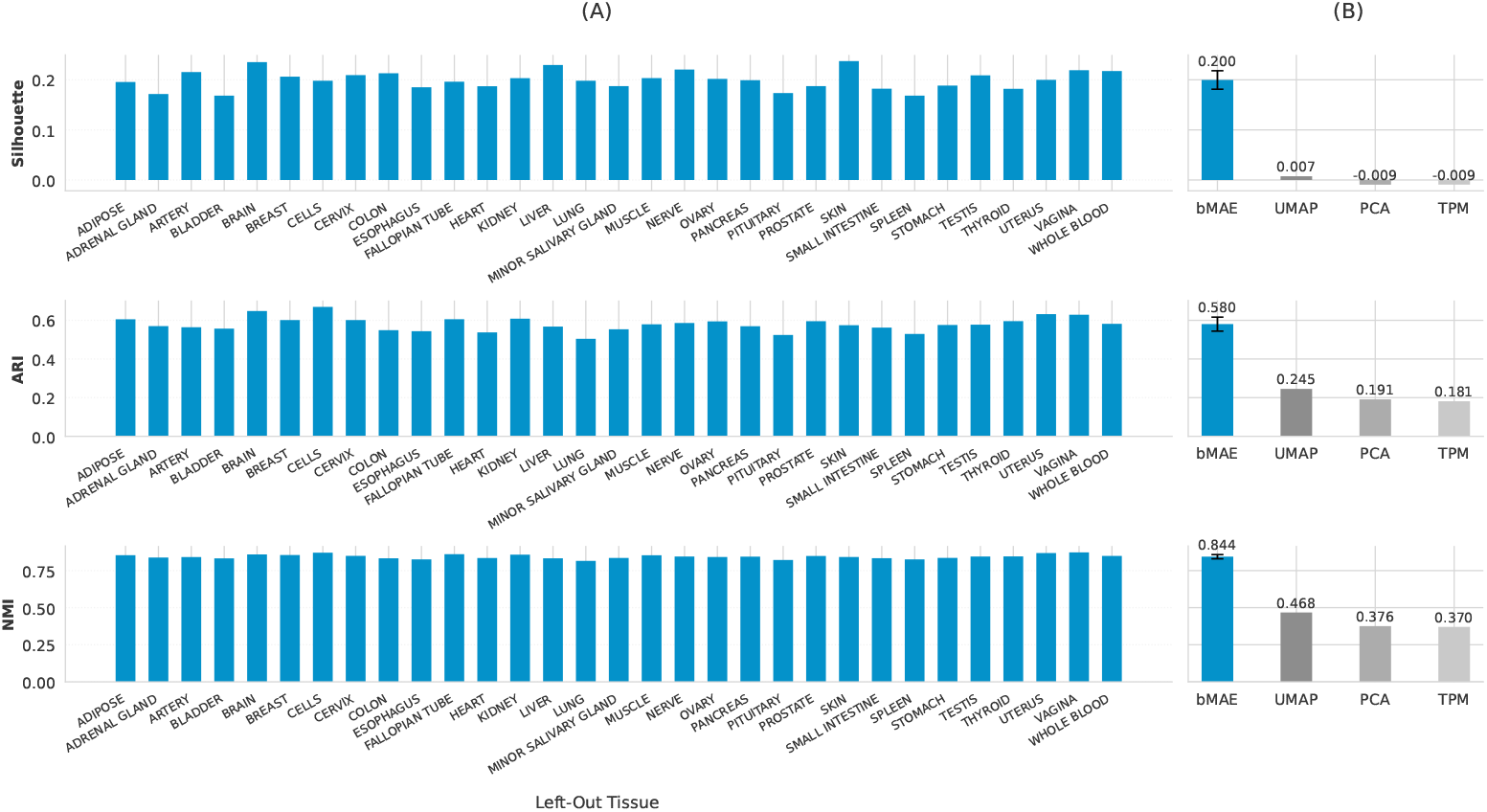
Latent Representation Comparison Among bMAE, UMAP, PCA, and TPM. (A): bMAE’s performance across the LOTO folds. (B): Comparison of the mean performance of bMAE versus commonly used dimension reductions

**Figure 4:**
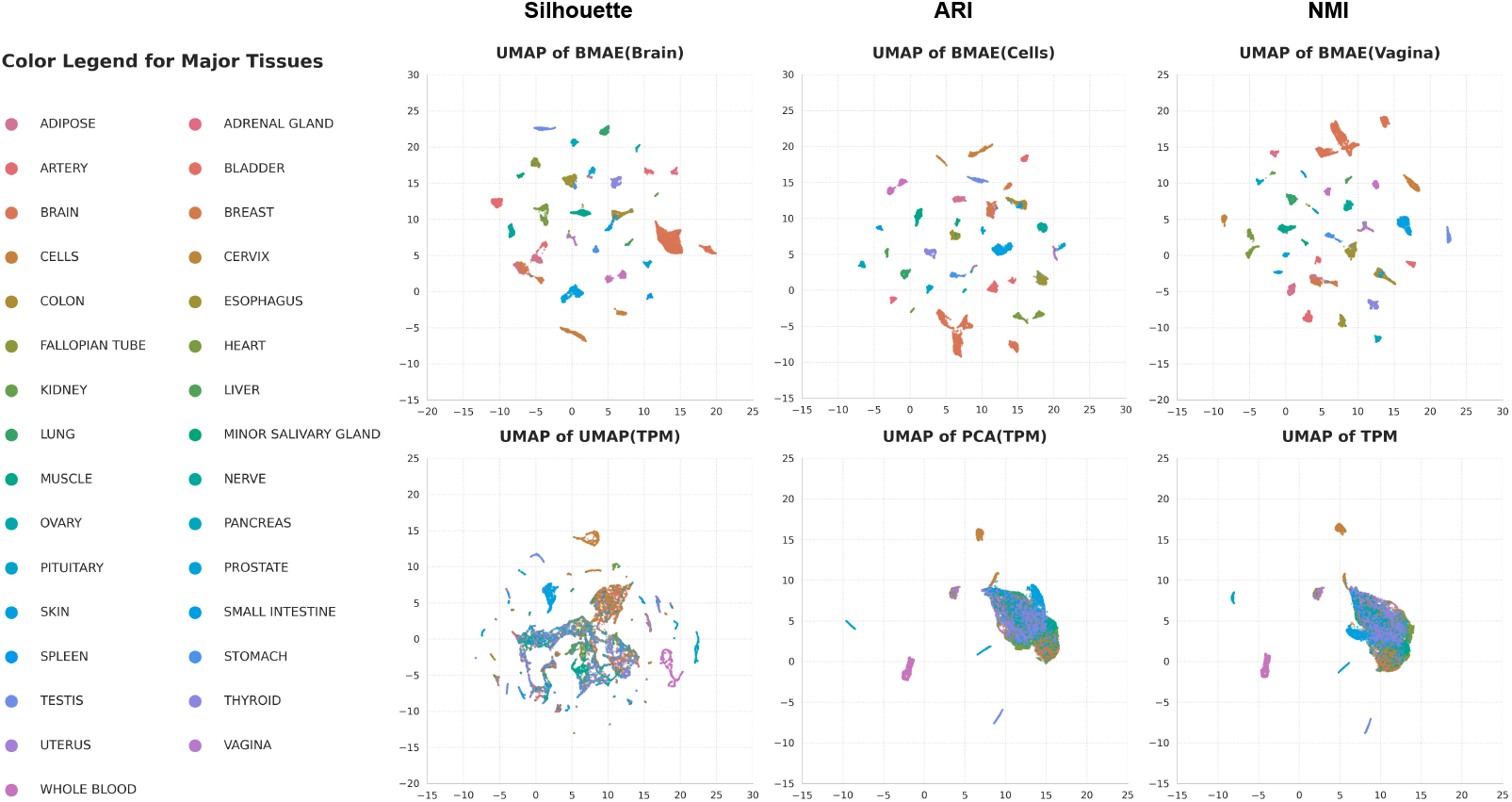
2-d UMAP of the 128-d Latent Representations of bMAE, UMAP, PCA, and TPM. For bMAE, the best for Silhoutte, ARI, and NMI, respectively, are shown.

Furthermore, Figures 5, 6, and 7 demonstrates that from the best case to the worst LOTO case, bMAE is able to distinguish the held out tissue from the rest of the tissues while UMAP mixes them despite seeing all of the tissues.

**Figure 5:**
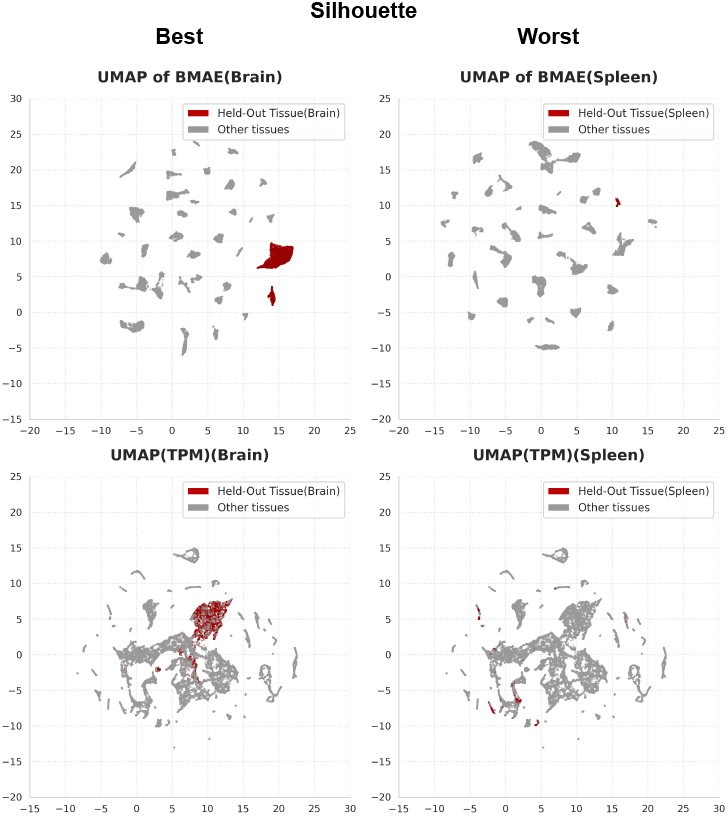
Best and Worst LOTO case for bMAE for Silhoutte compared to UMAP

**Figure 6:**
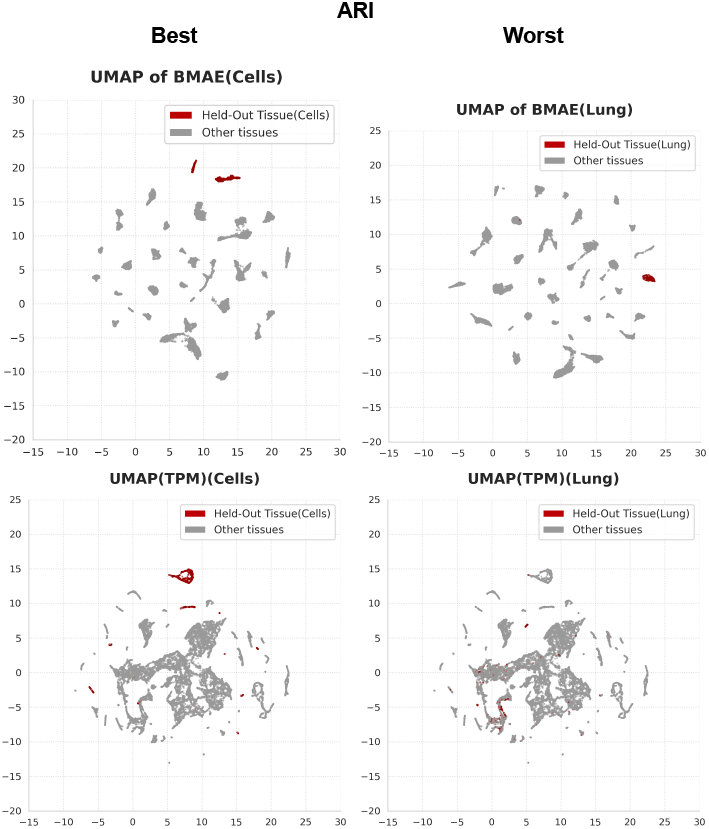
Best and Worst LOTO case for bMAE for ARI compared to UMAP

**Figure 7:**
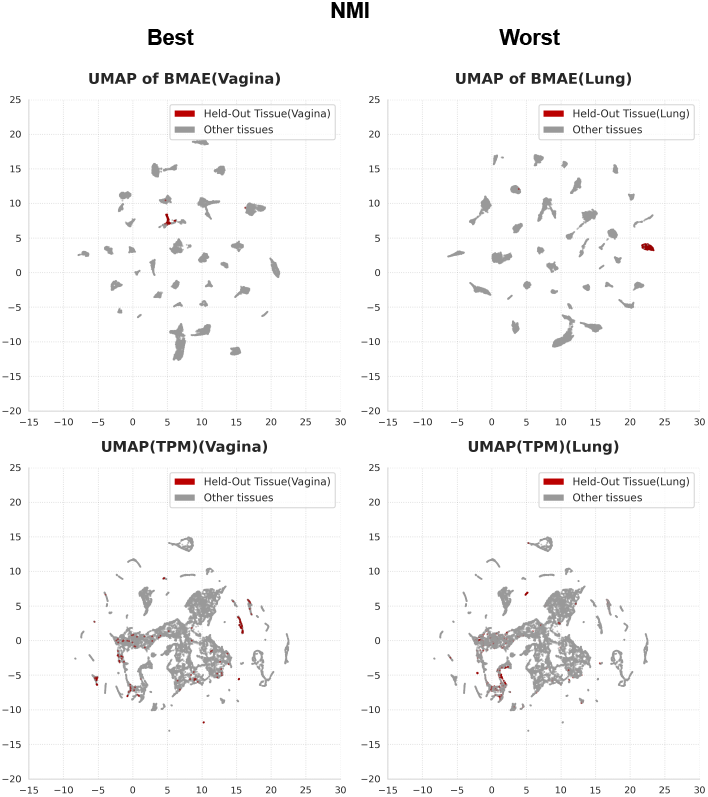
Best and Worst LOTO case for bMAE for NMI compared to UMAP

#### Subtissue Structure in Held-Out Tissues

To rigorously evaluate generalization to unseen tissue categories, we performed fine-grained analysis on the 10 major tissues containing multiple subtissues (e.g.,Brain with cortex, cerebellum, etc.; Artery with aorta, coronary, tibial). For each such tissue in LOTO cross-validation, we analyzed only the held-out tissue samples, using subtissue labels (rather than major tissue categories) as ground truth. Critically, the neural network models had never observed any samples from these major tissue categories during training, yet we evaluated whether they could discover the hierarchical subtissue structure within these completely unseen categories.

For this analysis, we compared only bMAE against classical methods (UMAP, PCA, Raw). Notably, PCA and UMAP were fitted on each held-out tissue’s samples with full access to that specific tissue’s expression patterns, providing them with a substantial advantage. In contrast, bMAE embeddings were generated by an encoder that had never seen this tissue category, relying entirely on learned features from the 30 other tissues.

Note that for because PCA dimensionality is constrained by *d* ≤ min(*n* − 1, *f*), where *n* is the number of samples and *f*is the number of features. Therefore, for held-out tissues with fewer than 128 samples, we reduced PCA dimensionality to match the sample count (*d* = *n*); otherwise, we used *d* = 128.

Despite the disadvantage, bMAE consistently outperformed all baselines at resolving subtissue structure (Table 2). Averaged across tissues with subtissue annotations, bMAE achieved silhouette 0.1616, ARI 0.3488, and NMI 0.3628, compared to UMAP’s 0.1498, 0.1948, 0.2007. Even when UMAP was fitted directly on the held-out tissue with full access to tissue-specific variance structure, bMAE’s zero-shot embeddings better separated subtissues in 7/10 tissues for silhouette, 9/10 tissues for ARI, and 10/10 tissues for NMI.

**Table 2:** Subtissue clustering performance on held-out major tissue categories containing multiple subtissues. Models evaluated only on samples from held-out tissue using subtissue labels. PCA and UMAP fitted on held-out tissue training samples; bMAE encoder never observed this tissue category during training. Values averaged across 10 tissues with subtissue annotations, reported as mean ± standard deviation.

| Method | Silhouette ↑ | ARI ↑ | NMI ↑ |
| --- | --- | --- | --- |
| **bMAE** | **0.1616 ± 0.18** | **0.3488 ± 0.31** | **0.3628 ± 0.29** |
| UMAP | 0.1498 | 0.1948 | 0.2007 |
| PCA | 0.0837 | 0.1163 | 0.1342 |
| Raw | 0.0837 | 0.1163 | 0.1342 |

This remarkable generalization demonstrates that bMAE learns transferable tissue-discriminative features rather than memorizing tissue-specific expression patterns.

### Ablation: No Masking, Fixed Masking, Variable Masking

In this section we show that the chosen variable masking rate during training helps the model achieve the performance or quality seen. To evaluate the contribution of variable masking schedules to representation learning, we conducted ablation studies comparing three model variants: (1) bMAE with variable masking, (2) bMAE-fixed trained with a constant mask rate of 0.85, and (3) AE, a standard autoencoder without masking during training. All models shared identical architecture and training configurations, differing only in their masking strategy.

#### Reconstruction and Recovery

We assessed each model’s ability to recover masked gene expression values using Pearson and Spearman correlation coefficients between predicted and true expression values. At test time, we applied a corruption rate of 0.85 to all models regardless of their training configuration, masking 85% of input genes and evaluating reconstruction quality on the masked subset. This evaluation was performed in a leave-one-tissue-out (LOTO) framework across all 31 tissue types, with each tissue serving as the held-out test set while the remaining 30 tissues comprised the training set.

#### Clustering Analysis for Major Tissues

As shown below, while Section 4.2.1 suggests that bMAE is not the best at reconstruction and recovery, it does, nevertheless, has better latent representations of the samples.

Comparing the three variants revealed a critical finding: while bMAE-fixed achieved superior reconstruction metrics (Section 4.2.1), bMAE with variable masking produced significantly higher-quality latent representations. Specifically, bMAE outperformed bMAE-fixed by 25.3% (31/31) in silhouette score, 3.1% (20/31) in ARI, and 2.2% (29/31) in NMI, while the standard AE without masking performed worst among neural approaches as shown in Table 4.

#### Statistical Testing

Shapiro-Wilk test was performed on the metrics across different LOTO cases. For the reconstruction metrics (Pearson and Spearman corelation), a *p* <= 0.05 was attained and hence treated as coming from a non-normal distribution. Table 3 summarizes the reconstruction performance averaged across all LOTO splits. For spearman correlation, no significant differences between any of them according to the Kruskal-Wallis H Test(*p* = 0.2249). As for pearson correlation, bMAE-fixed is statistically different with bMAE according to the Dunn Test (*p* = 0.0275).

**Table 3:** Reconstruction performance across ablation models. Metrics computed on masked genes (85% corruption) averaged over 31 LOTO splits. Values reported as mean ± standard deviation. Values in **bold** mean that they are the best. Multiple bold values mean for a column mean they have no statistical difference.

| Model | Pearson | Spearman |
| --- | --- | --- |
| bMAE-fixed | **0.9416 ± 0.117** | **0.9542 ± 0.018** |
| bMAE | 0.9267 ± 0.116 | **0.9561 ± 0.014** |
| AE | 0.9326 ± 0.123 | **0.9486 ± 0.018** |

For the latent representation quality metrics, the Shapiro-Wilk test, with a *p* >= 0.05, we treat silhouette, ARI, and NMI across LOTO cases as coming from a normal distribution and use the Levene Test and Bartlett Test to check the equality of variances. According to these tests, they do not have equal variances hence we use the Welch’s ANOVA test to check if they have statistical differences in their means. According to this test, there exists differences within each quality metric, and hence we use Games-Howell Test to find which model differs and for which quality metric. For Silhouette and NMI, they are all different. For ARI, bMAE-fixed and bMAE have no significant difference with *p* = 0.0707 > 0.05. Table 4 summarized the results for latent space quality metrics.

**Table 4:** Latent space quality metrics averaged across 31 LOTO splits. All methods project to 128 dimensions except Raw. Values in **bold** mean that they are the best. Multiple bold values for a column mean they have no statistical difference.

| Method | Silhouette ↑ | ARI ↑ | NMI ↑ |
| --- | --- | --- | --- |
| bMAE | **0.1996** $\pm 0.018$ | **0.5805** $\pm 0.036$ | **0.8438** $\pm 0.014$ |
| bMAE-fixed | 0.1593 $\pm 0.014$ | **0.5633** $\pm 0.022$ | 0.8265 $\pm 0.013$ |
| AE | -0.1023 $\pm 0.041$ | 0.2703 $\pm 0.068$ | 0.5372 $\pm 0.054$ |

#### Summary of Ablation Testing

From the statistical testing for several ablation cases, making the mask rate variable instead of fixed, does not affect the Spearman Correlation (monotonic trend) between the prediction and the true profile, however, it does affect the Pearson Correlation (linear trend) by lowering it. Despite this effect on the reconstruction, adding the variable masking schedule in the training, increases its performance on Silhouette (cohesion and separation) and NMI (mutual information between true labels and clustering); with no effect on ARI (agreement between true labels and clustering). Moreover, adding the masking in training, does not affect reconstruction metrics, but it does result in better latent representation.

This dissociation between reconstruction fidelity and representation quality demonstrates that the variable masking schedule acts as a powerful regularizer, forcing the encoder to extract robust features that generalize across different levels of data corruption. By encountering mask rates from 0.1 to 0.9 during training, bMAE learns representations that capture tissue identity at multiple scales of information availability, rather than overfitting to a single corruption pattern. This multiscale learning translates to embeddings that better preserve the biological signal, like tissue category membership, even when tested on entirely unseen tissue types in LOTO evaluation.

## Conclusions

### Generalization

We presented bMAE, a masked autoencoder framework with variable masking rate schedules for learning robust, generalizable representations from bulk tissue RNA-seq data. Our approach addresses a critical limitation in transcriptomic analysis: the inability of existing dimensionality reduction methods to capture tissue-discriminative features that transfer to entirely unseen tissue types.

Through rigorous leave-one-tissue-out cross-validation on the Adult GTEx dataset spanning 31 major tissue types, we demonstrated that bMAE substantially outperforms commonly used dimensionality reduction methods (PCA, UMAP) and raw expression profiles across multiple clustering quality metrics. Our learned 128-dimensional representations achieved mean silhouette scores of 0.1996, Adjusted Rand Index of 0.5805, and Normalized Mutual Information of 0.8438, representing 28.6-fold, 2.3-fold, and 1.8-fold improvements over UMAP respectively. Notably, bMAE embeddings revealed hierarchical subtissue structure within held-out major tissue categories, outperforming the other methods fitted directly on those tissues despite never observing them during training.

### Effectivity of Variable Masking Rate

Ablation studies revealed a key insight: variable masking schedules produce superior latent representations compared to fixed-rate masking, even though fixed-rate models achieve higher reconstruction accuracy. This demonstrates that point-wise prediction fidelity does not guarantee biologically meaningful embeddings. The variable schedule employed by bMAE acts as a powerful regularizer, forcing the encoder to extract robust, multi-scale features of tissue identity rather than memorizing tissue-specific expression patterns. The masking objective itself proved essential, as the standard autoencoder (AE) without masking produced substantially inferior representations with negative silhouette scores. This means that future work that uses masked autoencoding can benefit from applying this variable masking schedule for their training.

### Future Work

Several promising directions emerge from this work. First, extending bMAE to multi-modal integration, combining bulk RNA-seq with other high-throughput assays (e.g., ATAC-seq, proteomics, histology images) could yield richer tissue representations that capture regulatory and spatial context alongside transcriptional state. Second, investigating alternative masking strategies beyond uniform random masking, such as gene-module-aware masking based on known pathways or co-expression networks may better align the pretraining objective with biological structure. Third, exploring the learned latent space for biomarker discovery and interpretability through feature attribution methods could reveal which gene expression patterns drive tissue discrimination and whether these align with known biological markers.

Additionally, evaluating bMAE’s transfer learning capabilities across datasets from different sequencing platforms, laboratories, and disease contexts would establish its practical utility for cross-study integration. Extending the leave-one-tissue-out evaluation to leave-one-species-out settings could test whether the learned representations capture conserved mammalian tissue biology. Finally, adapting the framework for regression tasks, such as predicting continuous phenotypes (age, disease severity) or molecular traits (protein abundance, methylation levels) from latent representations would demonstrate broader applicability beyond tissue classification.

## Summary

This framework establishes that masked autoencoding with a variable corruption strategy can learn transferable biological principles from bulk RNA-seq data, enabling zero-shot tissue classification and few-shot learning in novel contexts. The success of bMAE highlights the untapped potential of self-supervised learning for bulk transcriptomics, a domain that has received less attention than single-cell methods despite bulk data’s continued practical advantages in cost, coverage, and clinical accessibility.

## Author contributions statement

Z.W. and M.Z.G.U. conceived and conducted the experiments, analysed the results, and wrote the manuscript. provided instructions and acquired funding. All authors reviewed and approved the final manuscript.

## Acknowledgments

This work was supported by Japan Society for the Promotion of Science Grant-in-Aid for Challenging Exploratory Research [JP25K22833]; and Japan Society for the Promotion of Science Research on Academic Transformation Areas (A) [JP22H05194].

## Data Availability

The dataset used can be accessed via the GTEx Portal, [Adult GTEx v10](https://storage.googleapis.com/adult-gtex/bulk-gex/v10/rna-seq/GTEx_Analysis_v10_RNASeQCv2.4.2_gene_tpm.gct.gz)

## Code Availability

Code can be found on Github [bmae](https://github.com/Mastopke304/bmae).

